# Gene expression imputation provides insight into the genetic architecture of frontotemporal dementia

**DOI:** 10.1101/2020.06.23.166355

**Authors:** Lianne M. Reus, Bogdan Pasaniuc, Danielle Posthuma, Toni Boltz, International FTD-Genomics Consortium (IFGC), Yolande A.L. Pijnenburg, Roel A Ophoff

**Author notes:** Correspondence to Lianne M. Reus. Postal address: De Boelelaan 1117, 1081 HV Amsterdam, The Netherlands.

## Abstract

The etiology of genetically sporadic frontotemporal dementia is poorly understood. Although genome-wide association studies for frontotemporal dementia have identified a small number of candidate risk regions, most of the risk genes remain largely unknown. To identify candidate genes with predicted expression levels associated with frontotemporal dementia, we integrated genome-wide summary statistics with external reference gene expression data, using a transcriptome-wide association studies approach.

FUSION software was used to leverage summary statistics on frontotemporal dementia (n=2,340 cases, n=7,252 controls) and clinical subtypes (behavioral variant frontotemporal dementia n=1,337 cases/2,754 controls; semantic dementia n=308 cases/616 controls; progressive non-fluent aphasia n=269 cases/538 controls, frontotemporal dementia with motor neuron disease n=200 cases/400 controls) from the International Frontotemporal Dementia Genomics Consortium with 53 expression quantitative loci tissue type panels (n=12,205 from five consortia). Significance was assessed using a 5% false discovery rate threshold. We identified 73 significant gene-tissue associations for frontotemporal dementia, representing 44 unique genes in 34 tissue types. Most significant findings were derived from dorsolateral prefrontal cortex splicing data (n=19 genes, 26%). Furthermore, the 17q21.31 inversion locus contained 23 significant associations, representing six unique genes whose predicted expression associated with frontotemporal dementia. Other top hits included *SEC22B* on chromosome 1, a gene involved in vesicle trafficking, *TRGV5* on chromosome 17 and *ZNF302* on chromosome 19. A single gene finding was observed for behavioral variant frontotemporal dementia (i.e., *RAB38* on chromosome 11) with evidence from multiple tissue types. For the other clinical subtypes no significant associations were observed.

We used transcriptome-wide association studies to prioritize candidate genes for frontotemporal dementia and identified a number of specific genes, including potential novel candidate genes (such as *SEC22B*) and previously reported risk regions (e.g., 17q.21.31). Most significant associations were observed in the dorsolateral prefrontal cortex, despite the modest sample size of the gene expression reference panel of this tissue type. This suggests that our findings are specific to frontotemporal dementia and are likely to be biologically relevant highlights of genes at different frontotemporal dementia risk loci that are contributing to the disease pathology.

## Introduction

Frontotemporal dementia (FTD) is a heterogeneous neurodegenerative disorder, characterized by frontal and/or temporal patterns of atrophy. Clinically, FTD patients present with the behavioral variant of FTD (bvFTD) or language variants, such as semantic dementia (SD) and progressive non-fluent aphasia (PNFA) (Rascovsky *et al*., 2011). In 10% of all cases, FTD co-occurs with motor neuron diseases (FTD-MND) (Seelaar *et al*., 2011).

Where FTD is mostly sporadic (80%), approximately 20% of all FTD cases are familial, with the most common Mendelian mutations including the hexanucleotide repeat expansion at the *C9ORF72* locus on chromosome 9, and mutations in microtubule-associated protein tau (*MAPT*) and progranulin (*GRN*) genes in and near the chromosome 17q21 inversion locus (Hutton *et al*., 1998; Baker *et al*., 2006; DeJesus-Hernandez *et al*., 2011; Renton *et al*., 2011; Greaves and Rohrer, 2019). Genome-wide association studies (GWAS) in FTD have also identified genetic risk variants, each having small associations with disease risk (Van Deerlin *et al*., 2010; Diekstra *et al*., 2014; Ferrari *et al*., 2014; Pottier *et al*., 2019). The number of known FTD disease susceptibility loci remains small due to limited power for discovery in the relatively small sample sizes of the GWAS studies thus far with n_cases_<5,000. At this time, it is poorly understood how genetic risk variants for FTD exert effects on etiology, while such knowledge is essential for understanding disease pathology and the development of therapeutic interventions.

Genetic risk variants identified in GWAS are often located in noncoding regions with and without regulatory motifs, outside the protein encoding sequences (Maurano *et al*., 2012). These risk variants are likely to predispose individuals to disease susceptibility by modulating mRNA expression levels, through local (*cis*) or distal (*trans*) expression quantitative trait loci (eQTL) (Nicolae *et al*., 2010). The FTD risk variant rs302652 nearby *RAB38* is a local eQTL, decreasing *RAB38* gene expression in monocytes (Ferrari *et al*., 2014) and potentially influencing bvFTD disease risk by modulating *RAB38* gene expression levels in specific brain areas. However, the joint effects of genetic risk loci for FTD on (differential) gene expression across multiple tissue types is unclear.

Transcriptome-wide association studies (TWAS) have emerged as a way to identify associations between traits and gene expression. The most common TWAS methods include PrediXcan, summary data–based Mendelian randomization (SMR) and FUSION (Gamazon *et al*., 2015; Gusev *et al*., 2016; Zhu *et al*., 2016). TWAS leverage the combined effects of multiple SNPs, either on individual-level (PrediXcan, SMR) or summary-level (s-PrediXcan, FUSION), on gene expression, thereby increasing power to find novel associations over a traditional GWAS when gene expression mediates risk (Gamazon *et al*., 2015; Gusev *et al*., 2016; Zhu *et al*., 2016). Imputation of the genetic control of gene expression is now widely used to decipher how GWAS identified alleles may contribute to disease risk and to identify specific candidate genes through which this effect is regulated. In this study, we performed a multi-tissue TWAS on sporadic FTD and its clinical subtypes, to identify genes whose changes in expression plays a role in FTD and to identify tissue types relevant to FTD. As a secondary aim of the study, we performed a TWAS-based enrichment analysis and explored whether FTD shows overlap in differential expression with psychiatric disorders that show clinical overlap with FTD.

## Materials and methods

### GWAS summary statistics

GWAS summary statistics from the International Frontotemporal Dementia Genomics Consortium (IFGC) (https://ifgcsite.wordpress.com/) on frontotemporal dementia (FTD; n=2,154 cases/4,308 controls) and FTD clinical subtypes, behavioral variant FTD (bvFTD; n=1,377 cases/2,754 controls), semantic dementia (SD; n=306 cases/616 controls), progressive non-fluent dementia (PNFA; n=269 cases/ 538 controls) and FTD with motor neuron disease (FTD-MND; n=200 cases/400 controls), were used (Table S1). Written informed consent was obtained from all participants according to the Declaration of Helsinki. For all study sites, the study was approved by the Medical Ethics Committee. Preprocessing and quality check procedures have been described previously (Ferrari *et al*., 2014). Single nucleotide polymorphisms (SNPs) were converted from chr:bp to rsID coordinates using Phase 3 1000 Genomes Project data (Genomes Project *et al*., 2015). Summary statistics were converted to LD-score format using the munge_stats.py utility from LDSC, leaving 1,068,995 SNPs for final analysis for all phenotypes (Bulik-Sullivan *et al*., 2015).

### Expression quantitative trait loci (eQTL) reference panels

Local eQTL datasets from five different cohorts (n=12,205) on 53 tissue types were downloaded from the FUSION website (http://gusevlab.org/projects/fusion) (Table 1). The five cohorts included the CommonMind Consortium (CMC, n=452) (Fromer *et al*., 2016), Netherlands Twin Registry (NTR, n=1,247) (Wright *et al*., 2014), The Cardiovascular Risk in Young Finns Study (YFS, n=1,264) (Laaksonen *et al*., 2017), Metabolic Syndrome in Men Study (METSIM, n=562) (Laakso *et al*., 2017) and the Genotype Tissue Expression project (GTEx) v7 (https://gtexportal.org/home/datasets, n=752). Local eQTLs were calculated by leveraging gene expression with genetic variation data (i.e., SNPs within ±1 Mb of the transcriptional start site (TSS) of the gene). More detailed information on genotyping and gene expression analyses for these datasets have been described previously: CMC (Gusev *et al*., 2018), NTR, YFS, METSIM (Gusev *et al*., 2016) and GTEx (Consortium., 2015).

**Table 1.**
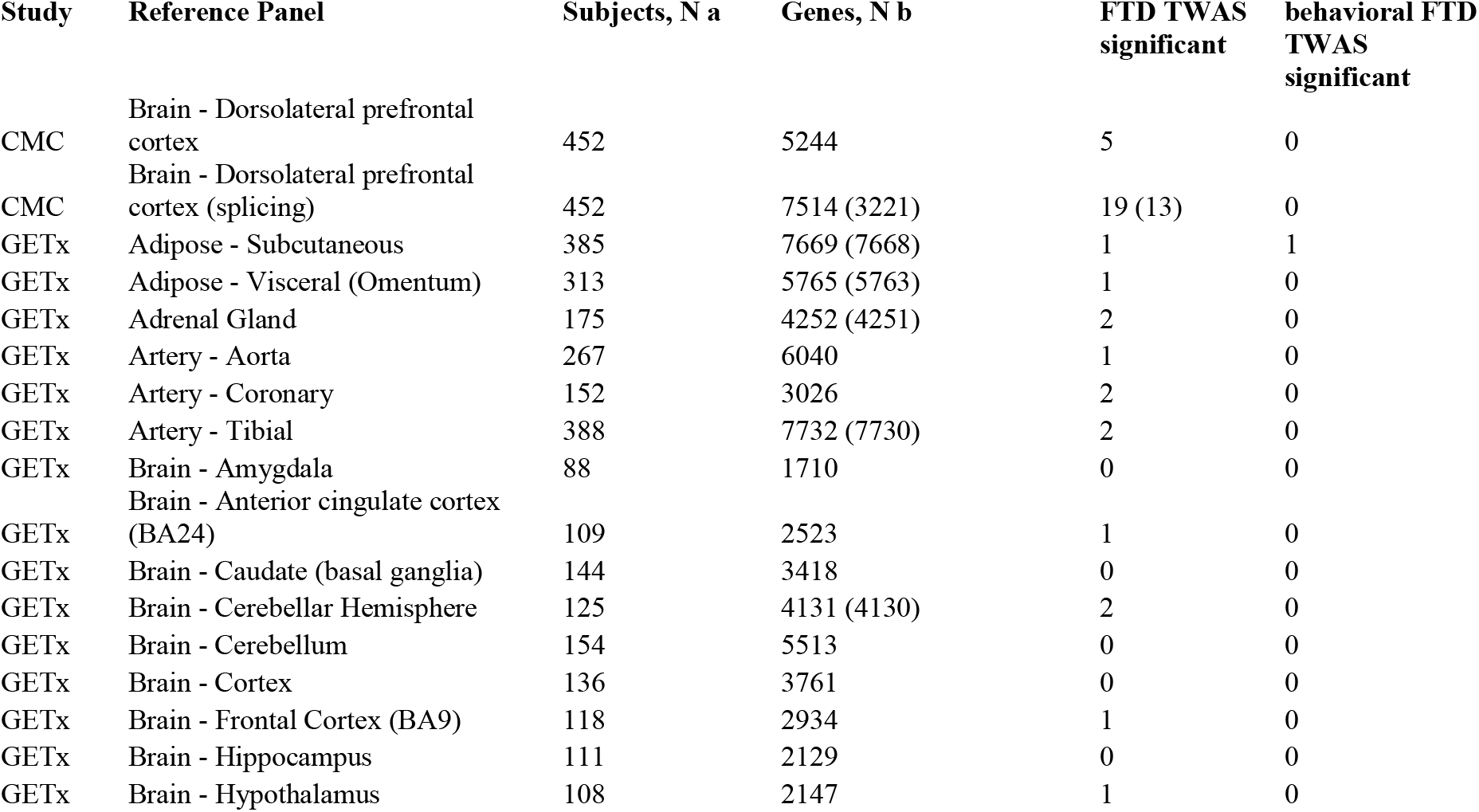
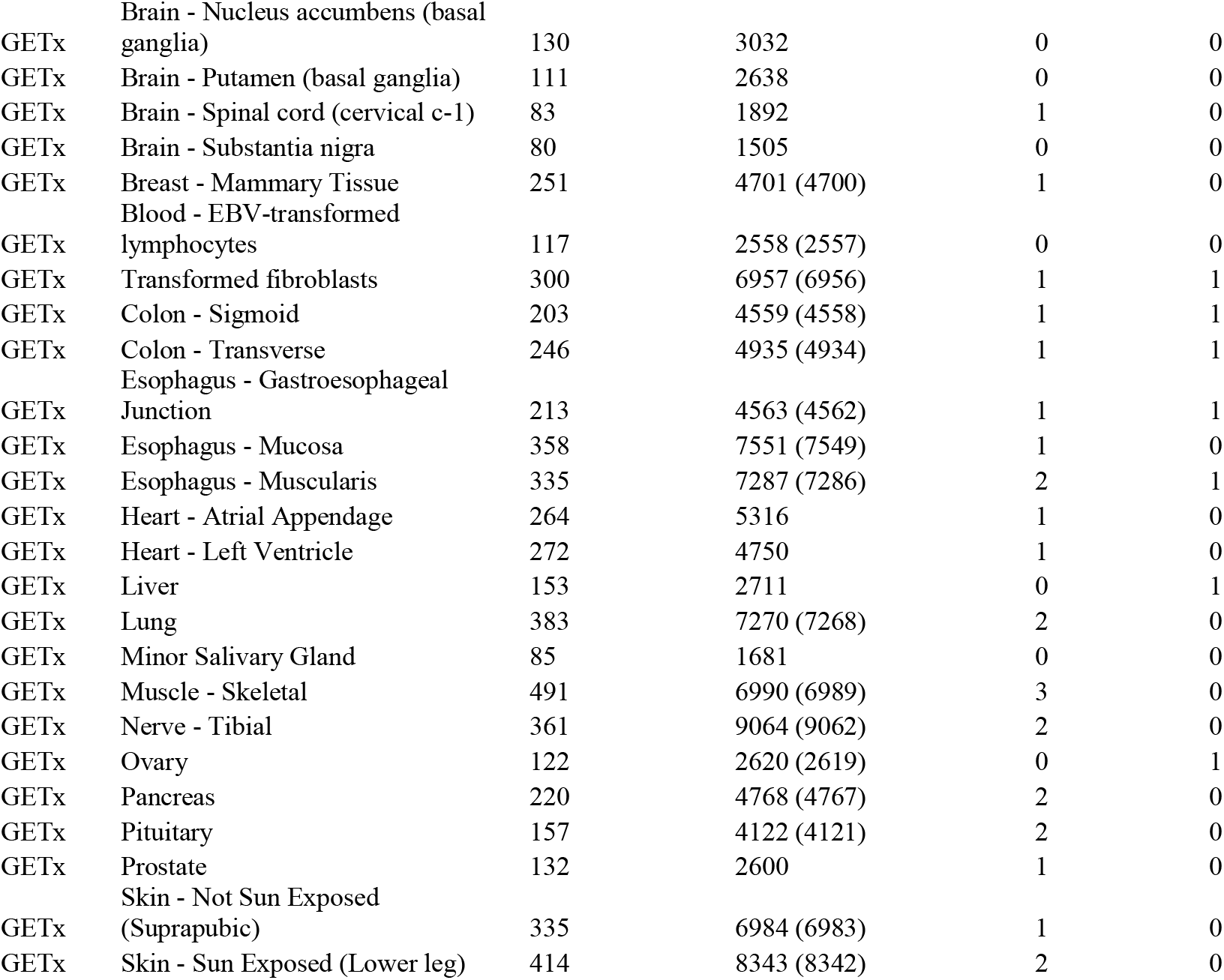
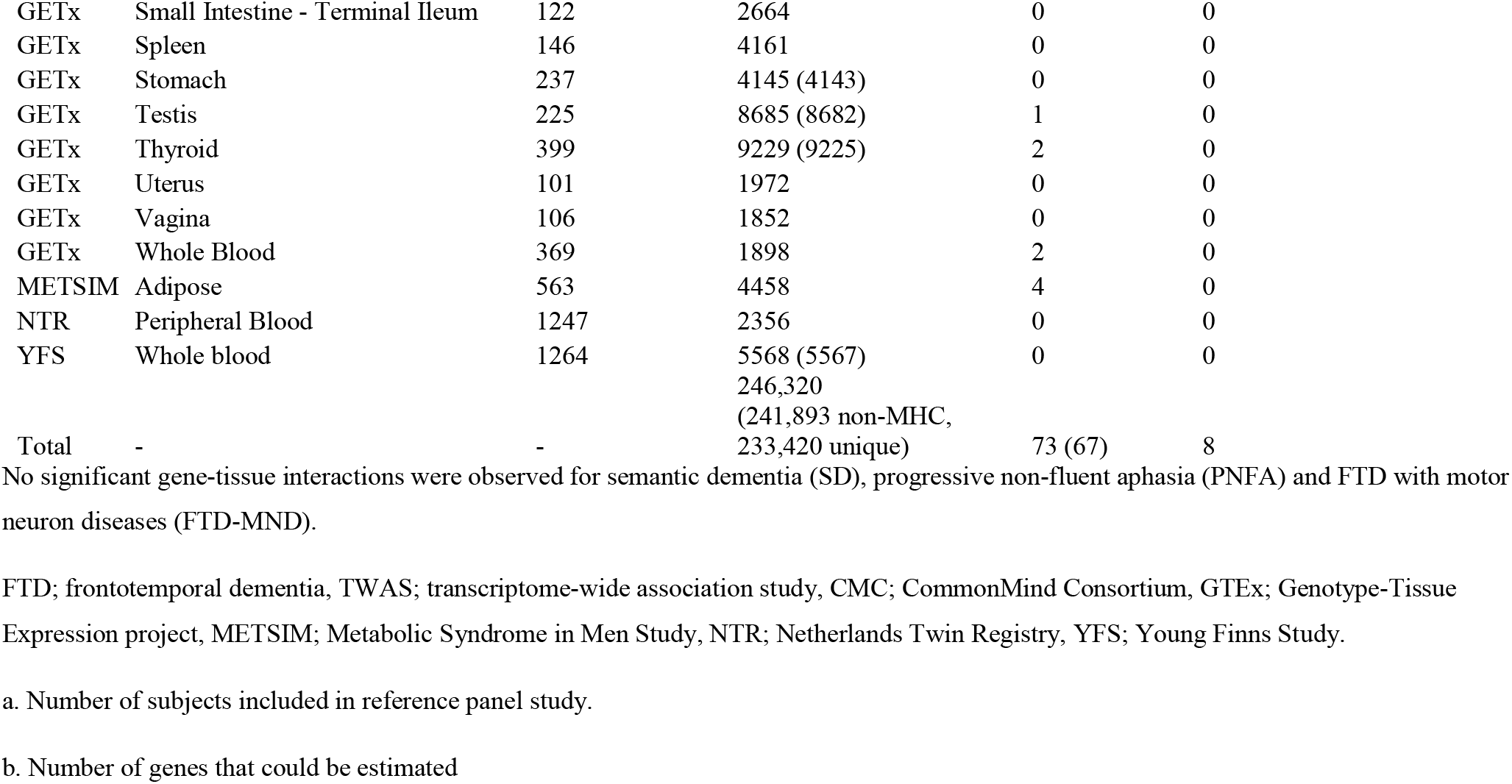
Descriptive statistics for tissue reference panels and TWAS results.

Local eQTL datasets from tissue types less relevant to FTD (e.g., blood) were included in this study, as local eQTLs are highly conserved across tissues (GTEx Consortium *et al*., 2017) and eQTL datasets with non-brain tissues consist of substantially larger sample sizes, thereby maximizing power to detect significant associations between local gene expression and FTD GWAS SNPs.

### FUMA

To examine the proportion of noncoding variants amongst FTD-risk SNPs, we annotated SNPs from the IFGC GWAS on FTD using Functional Mapping and Annotation (FUMA, https://fuma.ctglab.nl/) The most significant (p<5×10^− 6^) SNPs and SNPs in linkage disequilibrium (LD, r2≥0.6) with these were used for further inspection, using 1000 Genomes Project data (Genomes Project *et al*., 2015). Lead SNPs were defined as being independent from each other at r2>0.1. LD blocks of independent SNPs were merged into a genomic locus if they were closely located to each other (i.e., less than 250kb).

Lead and correlated SNPs were annotated for potential regulatory functions (RegulomeDB, RDB) (Boyle *et al*., 2012), 15-core chromatin state predicted by ChromHMM (Ernst and Kellis, 2012), functional consequences on gene functions annotated by ANNOVAR (Wang *et al*., 2010) and deleteriousness score (Combined Annotation Dependent Depletion, CADD) (Rentzsch *et al*., 2019).

### Statistical analysis

#### TWAS analysis

To identify genes whose local-regulated expression is associated with FTD and its clinical subtypes (i.e., bvFTD, SD, PNFA and FTD-MND), we performed TWAS analyses using FUSION software (http://gusevlab.org/projects/fusion/) (Gusev *et al*., 2016). FUSION estimates the genetic correlation between local gene expression and FTD, by integrating GWAS summary statistics with external gene expression reference panel data while accounting for LD structure among SNPs.

To study whether GWAS SNPs colocalized with eQTLs, we performed a Bayesian colocalization analysis for all associations with p_TWAS uncorrected_<0.05 using the COLOC package in R (https://cran.r-project.org/web/packages/coloc/) (Plagnol *et al*., 2009) implemented in FUSION. A joint analysis was performed to identify which genes are conditionally independent.

TWAS results are presented including the major histocompatibility (MHC) locus, as the FTD GWAS included genome-wide significant loci within the MHC region (Ferrari *et al*., 2014). Results on gene-tissue associations per phenotype (i.e., FTD, bvFTD, SD, PNFA and FTD-MND) were corrected for multiple comparisons using a 5% false discovery rate (FDR) significance threshold. Significant TWAS loci were identified as novel if the strongest FTD associated SNP was not nominal significant (*P*>0.05) in the IFGC GWAS (Ferrari *et al*., 2014) within ±1 Mb of the TSS of the gene’s region.

#### MESC analysis

In order to estimate the proportion of disease heritability mediated by local gene expression, we performed a Mediated Expression Score Regression (MESC) analysis per tissue type (https://github.com/douglasyao/mesc) (Yao et al., 2020). Here, we define h^2^_med_ as heritability mediated by local gene expression, h^2^_g_ as disease heritability and h^2^_med_/h^2^_g_ as the proportion of heritability mediated by local gene expression. First, for each gene, local heritability scores were estimated while accounting for LD structure. Genes were partitioned into bins according to their local heritability, as this has shown to provide unbiased h^2^_med_/h^2^_g_ estimates. Second, we estimated h^2^_med_/h^2^_g_ from expression scores estimated in the previous step and GWAS summary statistics on FTD. As MESC produces biased estimates for eQTL reference panels with small sample sizes, only eQTL datasets with sample size n>300 (n=17) were included.

#### Enrichment analysis

Competitive enrichment analysis on FTD TWAS results was performed using TWAS-based gene set enrichment analysis (TWAS-GSEA) (https://github.com/opain/TWAS-GSEA) (Pain *et al*., 2019). TWAS-GSEA is an adapted method of GWAS-based enrichment analysis implemented in software MAGMA (de Leeuw *et al*., 2015). In brief, this method examines whether TWAS results are enriched for specific pathways while accounting for LD structure. Per phenotype, TWAS-GSEA was performed simultaneously for all 53 eQTL datasets. If genes were present in multiple local eQTL datasets, the gene with the best prediction of expression (as estimated by cross-validated R^2^, MODELCV.R2) was used in the GSEA. Gene identifiers in TWAS result files were converted to Entrez ID format using the biomaRt package in R, resulting in 15,004 (14,813 non-MHC) unique Entrez IDs for FTD and all clinical FTD subtypes. TWAS results were tested for enrichment across 6,778 Gene Ontology (GO) biological processes gene sets. Per phenotype, results were corrected for the number of gene sets using a 5% FDR significance threshold.

### Data availability

Data can be made available upon request.

## Results

### Most risk variants for FTD are located in noncoding regions

For FTD, FUMA annotated 3,103 SNPs from thirteen independent lead SNPs located in ten genomic risk loci (Watanabe *et al*., 2017). Of all SNPs, 50.2% were located in intronic, 24.3% in non-coding RNA, 19.3% in intergenic, whereas only 1.4% was located in exonic regions. Most SNPs (93.1%) were located in open chromatin regions (range minimum chromatin state across 127 tissue/cell types=1-7) and 11.4% SNPs had potential regulatory elements, as indicated by a RDB score below 2 (Figure S1).

### Predicted gene expression levels show 73 associations with FTD

Predicted gene expression levels in 53 tissue types (range of genes per tissue type=1,505-9,229) were tested for association with FTD. We identified 73 significant gene-tissue associations for FTD, representing 44 (40 non-MHC) unique genes in 34 tissue types (Table 1, Table S2, Figure 1, Figure 2). The strongest genic FTD TWAS associations with supporting evidence from colocalization analyses included *ARL17B* on chromosome 17 (brain cerebellar hemisphere P_FDR_=9.02×10^− 22^), *ZNF302* on chromosome 19 (DLPFC splicing data P_FDR_=5.80×10^− 8^), *LRRC37A* (lung P_FDR_=1.58×10^− 5^), *SEC22B* on chromosome 1 (thyroid P_FDR_=2.28×10^− 3^) and *TRGV5P* on chromosome 17 (cells transformed fibroblasts P_FDR_= 2.39×10^− 3^) (Table 2, Table S3). Of all transcriptome-wide significant genes with supporting colocalization evidence, only the association of *SEC22B* with FTD was novel, showing no evidence for association in the FTD GWAS (minimal P within ±1Mb of the gene’s region=6.14×10^− 2^) (Ferrari *et al*., 2014) (Table S4).

**Figure 1.**
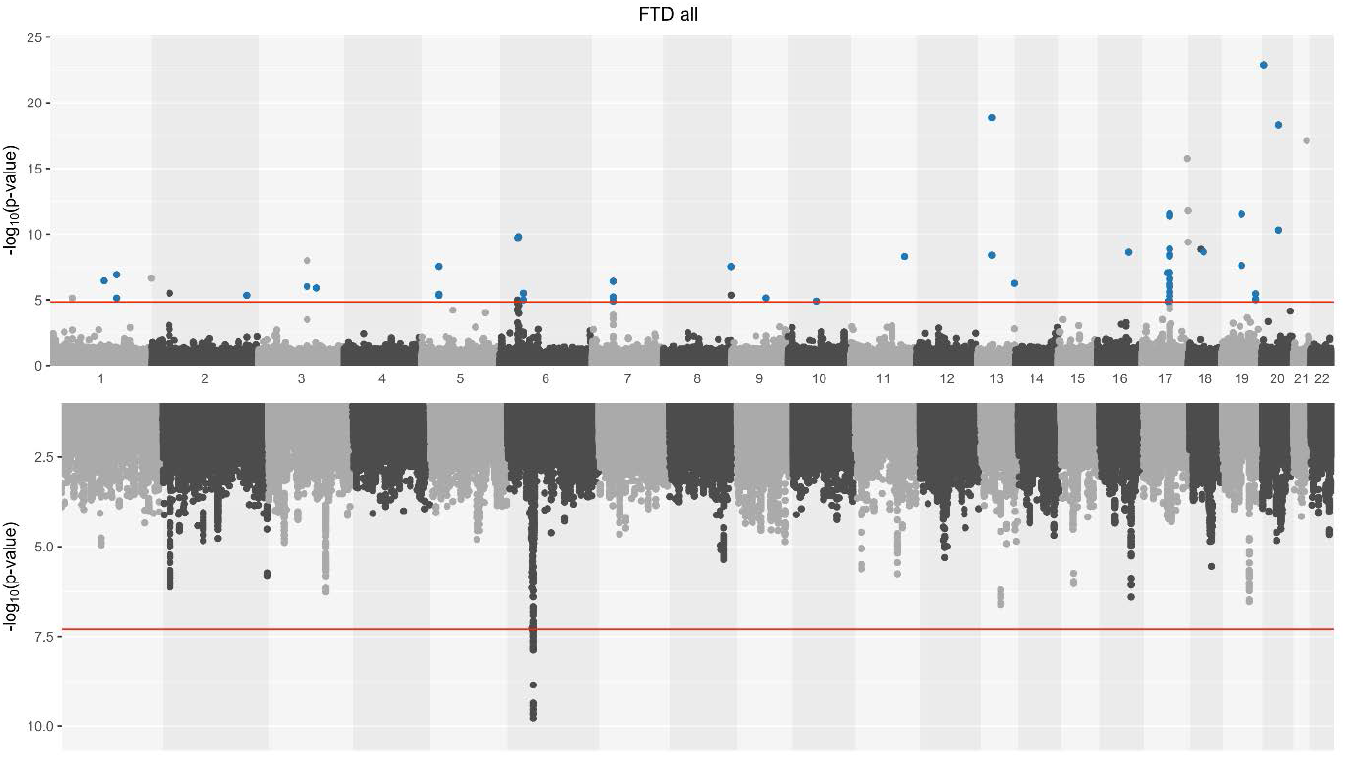
Manhattan plot on FTD TWAS (top) and GWAS (bottom). 44 unique genes were associated with FTD across 34 tissue types. Each point depicts a distinct gene-tissue association. TWAS hits with supporting evidence from colocalization analysis are highlighted blue. The red line depicts the significance threshold; *P*_FDR_<0.05 for TWAS and P<5e-8 for GWAS. FTD; frontotemporal dementia, TWAS; transcriptome-wide association study, GWAS: genome-wide association study.

**Figure 2.**
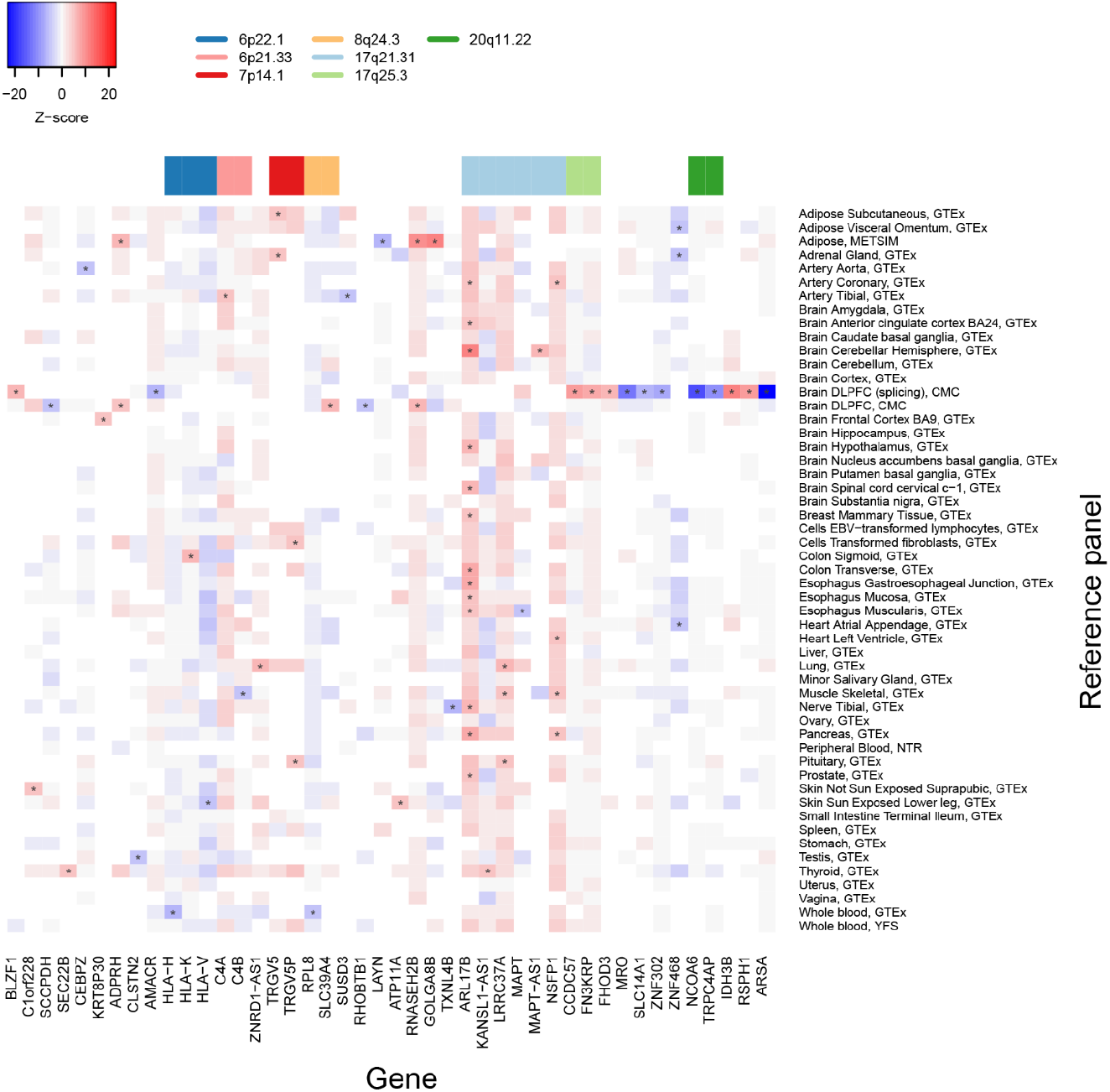
Heatmap of Z scores of genes with at least one transcriptome-wide significant association with FTD. Transcriptome-wide significant associations (*P*_FDR_<0.05) are depicted with an asterisk. Blank squares indicate that gene weights were not available in the reference panel. FTD; frontotemporal dementia, TWAS; transcriptome-wide association study, FDR; false-discovery rate.

**Table 2.** Transcriptome-wide significant associations with supporting evidence from colocalization analysis.

| **Location** | **Min p**<br>**(TWAS)** | **Min p**<br>**(GWAS)** | **Jointly significant** | **Marginally significant** |
| --- | --- | --- | --- | --- |
| 17.q21.31 | 1.83e-26 | 5.94e-05 | *ARL17B, LRRC37A, NSFP1* | *ARL17B KANSL1-AS1 LRRC37A NSFP1* |
| 13.q34 | 5.56e-07 | 2.30e-03 | *ATP11A* |  |
| 6.p21.33 | 1.00e-05 | 6.49e-04 | *C4A* |  |
| 3.q23 | 1.26e-06 | 3.07e-03 | *CLSTN2* |  |
| 2.q35 | 4.57e-06 | 9.48e-04 | *KRT8P30* |  |
| 10.q21.2 | 1.24e-05 | 2.69e-02 | *RHOBTB1* |  |
| 1.p12 | 3.61e-07 | 8.88e-02 | *SEC22B* |  |
| 9.q22.31 | 7.21e-06 | 6.14e-02 | *SUSD3* |  |
| 7.p14.1 | 3.89e-07 | 2.24e-02 | *TRGV5* | *TRGV5, TRGV5P* |
| 19.q13.11 | 3.06e-12 | 2.63e-02 | *ZNF302* | *ZNF302* |
| 19.q13.41 | 9.09e-06 | 6.04e-02 | *ZNF468* |  |

Min p (GWAS) represents the p value for the top SNP association within $\pm 1$ Mb of transcriptional site of the gene's region.
TWAS; transcriptome-wide association study, GWAS; genome-wide association study.

One region of interest is 17q21.31 on chromosome 17, which contained 23 significant associations, representing six unique genes (i.e., *ARL17B, KANSL1-AS1, LRRC37A, MAPT, MAPT-AS1* and *NSFP1*). This locus is an inversion polymorphism that has been associated previously with neurodegenerative tauopathies, but also with psychiatric disorders, such as autism spectrum disorders (Li *et al*., 2014; Pain *et al*., 2019). Gene expression of most gene-tissue pairs were highly correlated, except for *KANSL1-AS1*, *MAPT* and *MAPT-AS1* (Figure S2). For the majority of significant associations in 17q21.31 (n=16, 69.6%), colocalization supported model 4 (i.e., shared variant between gene expression and FTD, range PP3=0.001-0.22, range PP4=0.02-0.85).

Another region was 7p14.1, for which predicted gene expression of *TRGV5* and its pseudogene *TRGV5P* achieved transcriptome-wide significance in four different tissue types. Colocalization supported model 4 (range PP3=0.002-0.08, range PP4=0.76-0.98), suggesting that FTD and 17q21.31 gene expression share a single causal association.

### Most TWAS associations were detected in dorsolateral prefrontal cortex splicing data

The brain-derived reference panels contributed the most to the significant associations between gene expression and FTD (43.8%, 32 variants), with the majority derived from the dorsolateral prefrontal cortex (DLPFC) splicing data (19 variants, 13 unique, all outside MHC). The sample size and number of measured genes of the eQTL reference panel has shown to correlate with the number of significant hits (Gamazon *et al*., 2019). Despite the modest sample size (n_sample_=452) and number of measured genes (n_genes unique_=3,221, n_genes total_=7,514), the DLPFC splicing data accounted for 26% of all transcriptome-wide hits, thereby exceeding the number of significant hits compared to eQTL tissue types with larger sample sizes (e.g., 0% for YFS whole blood, n_sample_=1,264) and more measured genes (e.g., 3% for thyroid, n_genes, unique_=9,225, n_genes total_=9,229) (Figure S3, Figure S4).

MESC analysis showed that a substantial proportion of FTD heritability was mediated by the local component of gene expression levels (mean h^2^_med_=35(4.7)%). The tibial nerve had the highest heritability mediated by local gene expression levels (h^2^_med_=59.5(2.2)%), potentially reflecting a genetic component underlying the comorbidity underlying FTD and motor neuron diseases. For DLPFC splicing data the h^2^_med_ was 43.8(8.5)%, whereas for the eQTL panel with the largest sample size (YFS whole blood data) this was 12.6(7.4)%. A full overview of local mediated heritability is presented in Figure S5.

### Predicted gene expression levels on clinical subtypes separately show association with bvFTD only

Predicted gene expression levels in 53 tissue types (range of genes per tissue type=1,505-9,229) were tested for association with bvFTD, SD, PNFA and FTD-MND. Gene expression of *RAB38* on chromosome 11 was significantly associated with bvFTD risk in 8 out of 25 tissue panels (colon sigmoid P_FDR_=4.02×10^− 4^, range significant gene-tissue associations P_FDR_=4.02×10^− 4^-4.37×10^− 2^) (Figure 3, Figure S6, Table S5). Colocalization supported model 4 with a range posterior probability PP4 of 0.64-1.0 (range PP3=0.003-0.04) (Table S6). The former GWAS on bvFTD showed nominal evidence for the association of *RAB38* with FTD (rs302668 odd ratio(OR)=0.81(0.71-0.91); p_GWAS_=2.44×10^− 7^) (Ferrari *et al*., 2014). For SD, PNFA and FTD-MND, no significant transcriptome-wide associations were observed (Figure S7, S8, S9, Table S7-S12).

**Figure 3.**
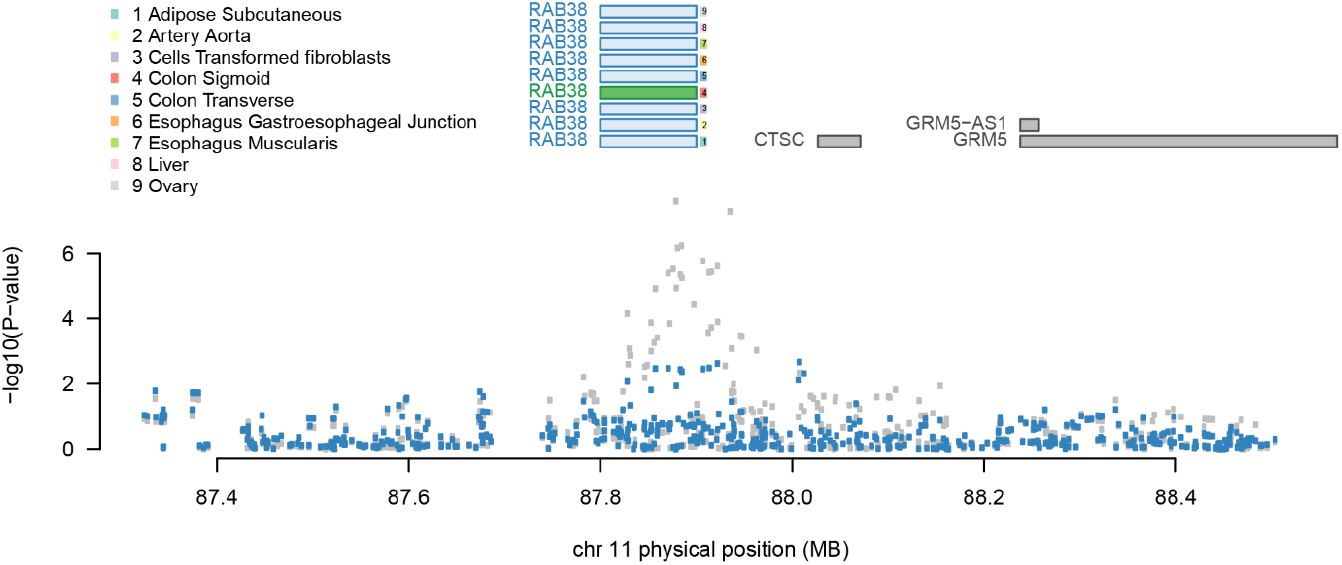
Regional association plot of *RAB38* for behavioral variant FTD. The top panel shows all of the genes in the locus. The marginally TWAS associated genes are highlighted in blue, and those that are jointly significant (i.e., *RAB38* in Colon Transverse) highlighted in green. The bottom panel shows a Manhattan plot of the GWAS data before (gray) and after (blue) conditioning on the green genes. This locus goes from being genome-wide significant to non-significant after conditioning on the predicted expression of *RAB38*. FTD; frontotemporal dementia.

### Implicated genes highlight involvement of amino acid transport in FTD pathogenesis

Full competitive results for the enrichment analysis on FTD and its clinical subtypes are presented in Table S13-S22. TWAS results for FTD were significantly enriched for ‘Sulfur amino acid transport’ (with MHC P_FDR_=0.04, without MHC P_FDR_=0.03) (Figure S11, S12). For all other gene sets and traits, no gene sets were significant after FDR correction.

### No genetic correlation between gene expression FTD and primary psychiatric disorders

Given the clinical similarities between FTD and primary psychiatric disorders, we also explored the genetic correlation between the predicted gene expression for FTD and schizophrenia, autism spectrum disorder and major depressive disorder, using RHOGE (Mancuso *et al*., 2017) (see Supplementary method section). No significant correlations were observed after FDR correction (Table S23,24, Figure S13).

## Discussion

In this study, we aimed to better understand the genetic etiology of sporadic FTD by identifying genes whose expression plays a role in FTD, using a TWAS approach with increased power of detecting loci compared to a traditional GWAS. We identified 73 significant gene-tissue associations for FTD, representing 44 unique genes in 34 tissue types. The 17q.21.31 inversion region was replicated as risk region for FTD. *SEC22B* was identified as likely novel risk gene for FTD. Interestingly, most associations were derived from DLPFC (a brain region relevant to FTD) splicing data, thereby providing some biological validation to the multi-tissue TWAS approach in FTD, and highlighting the importance of splicing events for disease risk (Li *et al*., 2016). Our results indicate that a large proportion of FTD risk loci modulate gene expression levels, and we highlight these genes as potential candidates for functional follow-up studies.

The majority of FTD risk variants were located in noncoding regions, demonstrating that these variants likely have regulatory functions. Forty-four genes were identified as differentially expressed in FTD. We replicated the 17q21.31 locus as risk factor for FTD. This region contained 23 significant associations from six different genes, including *ARL17B, KANSL1-AS1, LRRC37A, MAPT, MAPT-AS1* and *NSFP1*. The 17q21.31 region contains a common inversion polymorphism and has been associated with several neurodegenerative disorders (e.g., progressive supranuclear palsy, corticobasal degeneration, Alzheimer’s disease and FTD), but also with psychiatric disorders such as autism spectrum disorder (Myers *et al*., 2005; Webb *et al*., 2008; Li *et al*., 2014; Mishra *et al*., 2017; Gandal *et al*., 2018; Pain *et al*., 2019). Previous research has shown that different haplotypes of the 17q21.31 inversion affects expression of 17q21.31 genes in blood and different brain regions (de Jong *et al*., 2012). Here, we highlight the role of differential gene expression of 17q21.31 genes in the pathogenesis of FTD.

Another implicated gene was *SEC22B* on chromosome 1, which showed evidence for differential gene expression in FTD without achieving genome-wide significance in the corresponding FTD GWAS (P>0.05 within ± 1Mb of *SEC22B). SEC22B* is a vesicle trafficking protein, playing an important role in trafficking, autophagy and membrane fusion. The latter is essential for the development of the nervous system including axonal and dendritic growth (Petkovic *et al*., 2014). Little is known about the precise role of *SEC22B* in neurodegeneration, but differential expression of this gene in the brain has been associated with normal aging and Alzheimer’s disease (Berchtold *et al*., 2013; Zhao *et al*., 2016).

We found increased *C4A* gene expression to be significantly associated with FTD. The *C4* gene has two functionally different isoforms (i.e., C4A and C4B, both can vary in structure and copy number) and is located on the major histocompatibility (MHC) locus, a locus strongly associated with immune-related processes. Structural variation in *C4A/B* has been associated with schizophrenia, probably affecting synaptic pruning (Sekar *et al*., 2016; Kamitaki *et al*., 2020). The potential role of *C4* (structure) in the etiology of FTD has not been fully understood yet. Human post-mortem and mice model studies on FTD demonstrate an association between upregulated *C4A* gene expression and aggregation of transactive response (TAR) DNA binding protein, 43 kDa (TDP), one of the most common pathological subtypes of frontotemporal lobar degeneration (FTLD) (Chen-Plotkin *et al*., 2008; Wu *et al*., 2019). Upregulated *C4A* gene expression is probably generic to neurological disorders rather than specific for FTD, as it has been observed in other neurological and psychiatric disorders (McCarthy *et al*., 2019).

Proteins differentially expressed in FTD showed enrichment for the transport of sulfur amino acids. Sulfur amino acids (e.g., methionine and cysteine) are sensitive to oxidative modifications by reactive oxygen-containing species (ROS). The transport of sulfur amino acids is essential for the synthesis of antioxidants. For example, transport of L-cystine (i.e., oxidized form of cysteine) is needed for the production of antioxidant glutathione in the brain (McBean and Flynn, 2001). A balance between the production of ROS and antioxidants protects cells against invaders. However, an imbalance leads to increased oxidative stress, which is particularly damaging to cells in high demand of oxygen, such as neuronal cells (Haque *et al*., 2019). Increased oxidative stress has been associated with aging, and has been observed in several disorders, including FTD (Stadtman *et al*., 2005; Palluzzi *et al*., 2017; Haque *et al*., 2019).

The DLPFC contributed toward a large number of significant transcriptome-wide findings, thereby highlighting the topology-specific neurodegenerative nature of FTD. MESC analysis, an approach to examine the genome-wide distribution of heritability, showed that the tibial nerve had the largest proportion of heritability mediated by local gene expression, which may reflect the comorbidity of FTD with motor neuron diseases. We also observed various associations outside the brain, potentially highlighting the importance of other organ systems in FTD. In line with this, other organ systems, such as the gastrointestinal and musculoskeletal system, have been associated with FTD (Ikegami *et al*., 2000; Ahmed *et al*., 2016). On the other hand, we included local eQTL data from many tissue types - also those that seemingly are less disease-relevant - to increase power and to include as many genes in this exploratory study. As a result we may not have detected the true mechanism of disease due to a shared cross-tissue regulatory architecture of eQTLs between the tissue types related and non-related to FTD (GTEx Consortium *et al*., 2017; Wainberg *et al*., 2019). This is illustrated by our finding on bvFTD, for which we only identified differential regulatory gene expression of *RAB38* in tissue types outside the brain. As *RAB38* is expressed throughout the brain (https://gtexportal.org/home/gene/RAB38) but not available in the brain tissue panels we used, we hypothesize that differential expression of *RAB38* in the brain contributes to bvFTD disease risk too.

This study is a starting point for bridging the gap between genetic variation and disease pathogenesis involving specific genes in FTD. Nevertheless, several limitations should be taken into account. First, where TWAS increases power over a traditional GWAS study, the small sample size of the current FTD GWAS study still reduces the power to find novel transcriptome-wide associations. A second major limitation is that this study does not address the pathological heterogeneity in FTD. The most common pathological subtypes of FTD include abnormal aggregation of tau (FTLD-tau) and FTLD-TDP (Mackenzie *et al*., 2010). As we performed a TWAS on the clinical entity of FTD, this study provides only insights into generic mechanisms underlying FTD but not into specific mechanisms underlying pathological subtypes. Additional studies are required to gain more insight into distinct mechanisms underlying pathological subtypes of FTD. Finally, it should be noted that TWAS or colocalization analysis cannot be used for causal inference (Wainberg *et al*., 2019). It is therefore essential that our efforts will be extended to functional validation, to further understand the relationship between FTD and genes reported in this study.

Results presented in this study could be used as a point of reference in future genetic association studies on FTD. We provide evidence for the contribution of many genes to the pathogenesis of FTD, including potential novel (i.e., *SEC22B*) and previously reported FTD risk loci (e.g., 17q21.31 inversion region). Most associations were detected in DLPFC splicing data, but tissues outside the brain may be involved in FTD as well. However, functional validation is needed as TWAS is sensitive to detecting associations not relevant for disease if the disease-relevant tissue is not well-represented across reference panels. Identifying which biological processes are genetically influenced by FTD is important for understanding the disease etiology, and eventually for the development of treatments.

## Acknowledgement(s)

We thank and acknowledge the International FTD-Genomics Consortium (IFGC).

## List of abbreviations

bvFTD: behavioral variant of frontotemporal dementia
CADD: Combined Annotation Dependent Depletion
CMC: CommonMind Consortium
DLPFC: dorsolateral prefrontal cortex
eQTL: expression quantitative trait loci
FDR: false discovery rate
FTD: frontotemporal dementia
FTD-MND: frontotemporal dementia with motor neuron diseases
FTLD: frontotemporal lobar degeneration
FTLD-tau: frontotemporal lobar degeneration with tau pathology frontotemporal lobar degeneration with transactive response DNA binding
FTLD-TDP: protein pathology
FUMA: Functional Mapping and Annotation
GO: Gene Ontology
GTEx: Genotype-Tissue Expression project
GWAS: genome-wide association studies
IFGC: International Frontotemporal Dementia Genomics Consortium
LD: linkage disequilibrium
MESC: Mediated Expression Score Regression
METSIM: Metabolic Syndrome in Men Study
MHC: major histocompatibility
NTR: Netherlands Twin Registry
PNFA: progressive non-fluent aphasia
PP: posterior probability
ROS: reactive oxygen-containing species
SD: semantic dementia
SMR: summary data–based Mendelian randomization
SNPs: single nucleotide polymorphisms
TDP: transactive response DNA binding protein
TWAS: transcriptome-wide association study
TWAS-GSEA: TWAS-based gene set enrichment analysis
YFS: The Cardiovascular Risk in Young Finns Study

