## Supplemental information for "Gene expression imputation provides insight into the genetic architecture of frontotemporal dementia"

### Supplementary materials - Multi-tissue gene expression imputation provides insight into the genetic architecture of frontotemporal dementia

#### Supplementary table captions

**Table S1. Characteristics GWAS summary statistics.**

GWAS; genome-wide association studies, FTD; frontotemporal dementia, bvFTD; behavioural variant FTD, SD; semantic dementia, PNFA; progressive non-fluent aphasia, FTD-MND; frontotemporal dementia with motor neuron disease.

**Table S2. FTD TWAS results.**

FTD; frontotemporal dementia, TWAS; transcriptome-wide association study, CHR; chromosome, P0; gene start, P1; gene end, HSQ; heritability of the gene, BEST.GWAS.ID; rsID of the most significant SNP in locus, BEST.GWAS.Z; Z-score of the most significant GWAS SNP in locus, EQTL.ID; rsID of the best eQTL in the locus, EQTL.R2; cross-validation R2 of the best eQTL in the locus, EQTL.Z; Z-score of the best eQTL in the locus, EQTL.GWAS.Z; GWAS Z-score for this eQTL, NSNP; number of SNPs in the locus, MODELCV.R2; cross-validation R2 of the best performing model, MODELCV.PV; cross-validation P-value of the best performing model, SNP; single nucleotide polymorphism, FDR; false-discovery rate, eQTL; expression quantitative trait locus. Only TWAS associations with $p_{\text{TWAS uncorected}}<0.05$ are presented in the table.

**Table S3. FTD colocalization results.**

FTD; frontotemporal dementia, TWAS; transcriptome-wide association study, COLOC; colocalization, PP; posterior probability.

**Table S4. Significance in former FTD GWAS (Ferrari *et al.*, 2014) of transcriptome-wide significant genes with supporting colocalization evidence**. Min p (GWAS) represents the p value for the top SNP association within ±1Mb of transcriptional start site (TSS) of the gene's region.

GWAS; genome-wide association study.

**Table S5. BvFTD TWAS results.**

bvFTD; behavioral variant frontotemporal dementia, TWAS; transcriptome-wide association study, CHR; chromosome, P0; gene start, P1; gene end, HSQ; heritability of the gene,

BEST.GWAS.ID; rsID of the most significant SNP in locus, BEST.GWAS.Z; Z-score of the most significant GWAS SNP in locus, EQTL.ID; rsID of the best eQTL in the locus, EQTL.R2; cross-validation R2 of the best eQTL in the locus, EQTL.Z; Z-score of the best eQTL in the locus, EQTL.GWAS.Z; GWAS Z-score for this eQTL, NSNP; number of SNPs in the locus, MODELCV.R2; cross-validation R2 of the best performing model, MODELCV.PV; cross-validation P-value of the best performing model, SNP; single nucleotide polymorphism, FDR; false-discovery rate, eQTL; expression quantitative trait locus. Only TWAS associations with $p_{\text{TWAS uncorected}}$<0.05 are presented in the table.

**Table S6. BvFTD colocalization results.**

bvFTD; behavioral variant frontotemporal dementia, TWAS; transcriptome-wide association study, COLOC; colocalization, PP; posterior probability.

**Table S7. SD TWAS results.**

SD; semantic dementia, TWAS; transcriptome-wide association study, CHR; chromosome, P0; gene start, P1; gene end, HSQ; heritability of the gene, BEST.GWAS.ID; rsID of the most significant SNP in locus, BEST.GWAS.Z; Z-score of the most significant GWAS SNP in locus, EQTL.ID; rsID of the best eQTL in the locus, EQTL.R2; cross-validation R2 of the best eQTL in the locus, EQTL.Z; Z-score of the best eQTL in the locus, EQTL.GWAS.Z; GWAS Z-score for this eQTL, NSNP; number of SNPs in the locus, MODELCV.R2; cross-validation R2 of the best performing model, MODELCV.PV; cross-validation P-value of the best performing model, SNP; single nucleotide polymorphism, FDR; false-discovery rate, eQTL; expression quantitative trait locus. Only TWAS associations with $p_{\text{TWAS uncorected}}$<0.05 are presented in the table.

**Table S8. SD colocalization results.**

SD; semantic dementia, TWAS; transcriptome-wide association study, COLOC; colocalization, PP; posterior probability.

**Table S9. PNFA TWAS results.**

PNFA; progressive non-fluent aphasia, TWAS; transcriptome-wide association study, CHR; chromosome, P0; gene start, P1; gene end, HSQ; heritability of the gene, BEST.GWAS.ID; rsID of the most significant SNP in locus, BEST.GWAS.Z; Z-score of the most significant GWAS SNP in locus, EQTL.ID; rsID of the best eQTL in the locus, EQTL.R2; cross-validation R2 of the best eQTL in the locus, EQTL.Z; Z-score of the best eQTL in the locus,

EQTL.GWAS.Z; GWAS Z-score for this eQTL, NSNP; number of SNPs in the locus, MODELCV.R2; cross-validation R2 of the best performing model, MODELCV.PV; cross-validation P-value of the best performing model, SNP; single nucleotide polymorphism, FDR; false-discovery rate, eQTL; expression quantitative trait locus. Only TWAS associations with $p_{TWAS\ uncorected}<0.05$ are presented in the table.

**Table S10. PNFA colocalization results.**
PNFA; progressive non-fluent aphasia, TWAS; transcriptome-wide association study, COLOC; colocalization, PP; posterior probability.

**Table S11. FTD-MND TWAS results.**
FTD-MND; frontotemporal dementia with motor neuron disease, TWAS; transcriptome-wide association study, CHR; chromosome, P0; gene start, P1; gene end, HSQ; heritability of the gene, BEST.GWAS.ID; rsID of the most significant SNP in locus, BEST.GWAS.Z; Z-score of the most significant GWAS SNP in locus, EQTL.ID; rsID of the best eQTL in the locus, EQTL.R2; cross-validation R2 of the best eQTL in the locus, EQTL.Z; Z-score of the best eQTL in the locus, EQTL.GWAS.Z; GWAS Z-score for this eQTL, NSNP; number of SNPs in the locus, MODELCV.R2; cross-validation R2 of the best performing model, MODELCV.PV; cross-validation P-value of the best performing model, SNP; single nucleotide polymorphism, FDR; false-discovery rate, eQTL; expression quantitative trait locus. Only TWAS associations with $p_{TWAS\ uncorected}<0.05$ are presented in the table.

**Table S12. FTD-MND colocalization results.**
FTD-MND; frontotemporal dementia with motor neuron disease, TWAS; transcriptome-wide association study, COLOC; colocalization, PP; posterior probability.

**Table S13-S22**. Competitive TWAS-GSEA results for FTD, bvFTD, SD, PNFA and FTD-MND.

**Table S23. Genome-wide genetic correlation between FTD and psychiatric disorders as a function of predicted gene expression excluding the MHC region.**
FTD; frontotemporal dementia, bvFTD; behavioral variant FTD, SD; semantic dementia, PNFA; progressive non-fluent aphasia, FTD-MND; FTD with motor neuron disease, SCZ; schizophrenia, MDD; major depressive disorder, RHOGE; rho gene expression, TSTAT; t-statistics, DF; degrees of freedom, FDR; false-discovery rate.

**Table S24. Genome-wide genetic correlation between FTD and psychiatric disorders as a function of predicted gene expression including the MHC region.**

FTD; frontotemporal dementia, bvFTD; behavioral variant FTD, SD; semantic dementia, PNFA; progressive non-fluent aphasia, FTD-MND; FTD with motor neuron disease, SCZ; schizophrenia, MDD; major depressive disorder, RHOGE; rho gene expression, TSTAT; t-statistics, DF; degrees of freedom, FDR; false-discovery rate.

**Table S25. Characteristics cis expression quantitative loci (eQTL) datasets.**

CMC; CommonMind Consortium, GTEx; Genotype-Tissue Expression project, METSIM; Metabolic Syndrome in Men Study, NTR; Netherlands Twin Registry, YFS; Young Finns Study.

#### Supplementary figures

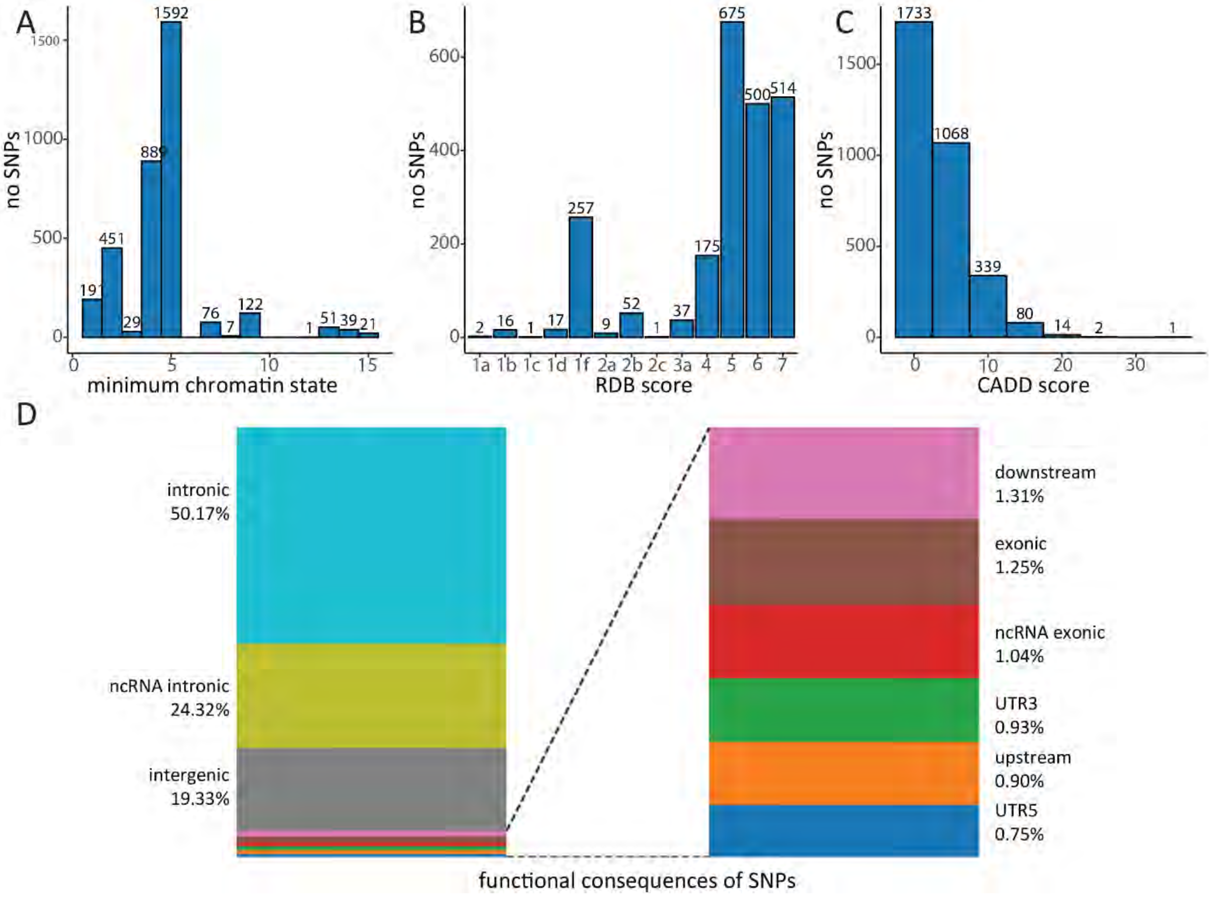


**Figure S1. Functional consequences of SNPs in FTD GWAS.**

FTD; frontotemporal dementia, GWAS; genome-wide association studies, SNPs; single nucleotide polymorphisms, RDB; RegulomeDB, CADD; Combined Annotation Dependent Depletion, ncRNA; non-coding ribonucleic acid, UTR3; three prime untranslated region, UTR5; three prime untranslated region.

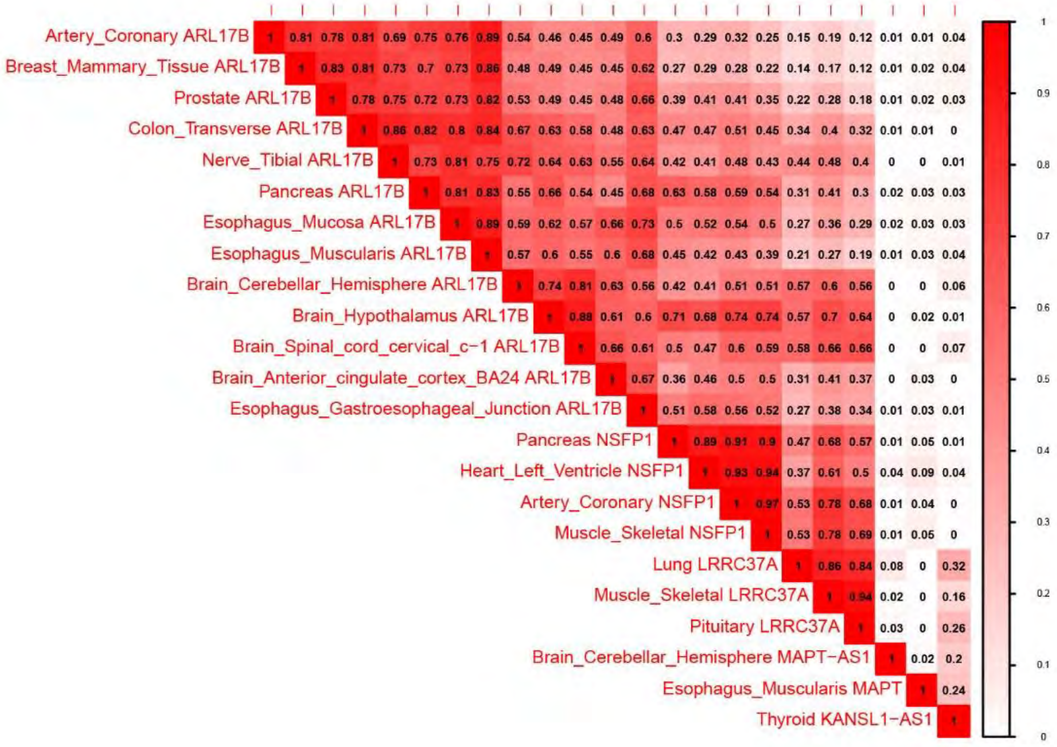


**Figure S2. Correlations between transcriptome-wide gene-tissue features on chromosome 17q.21.31.**

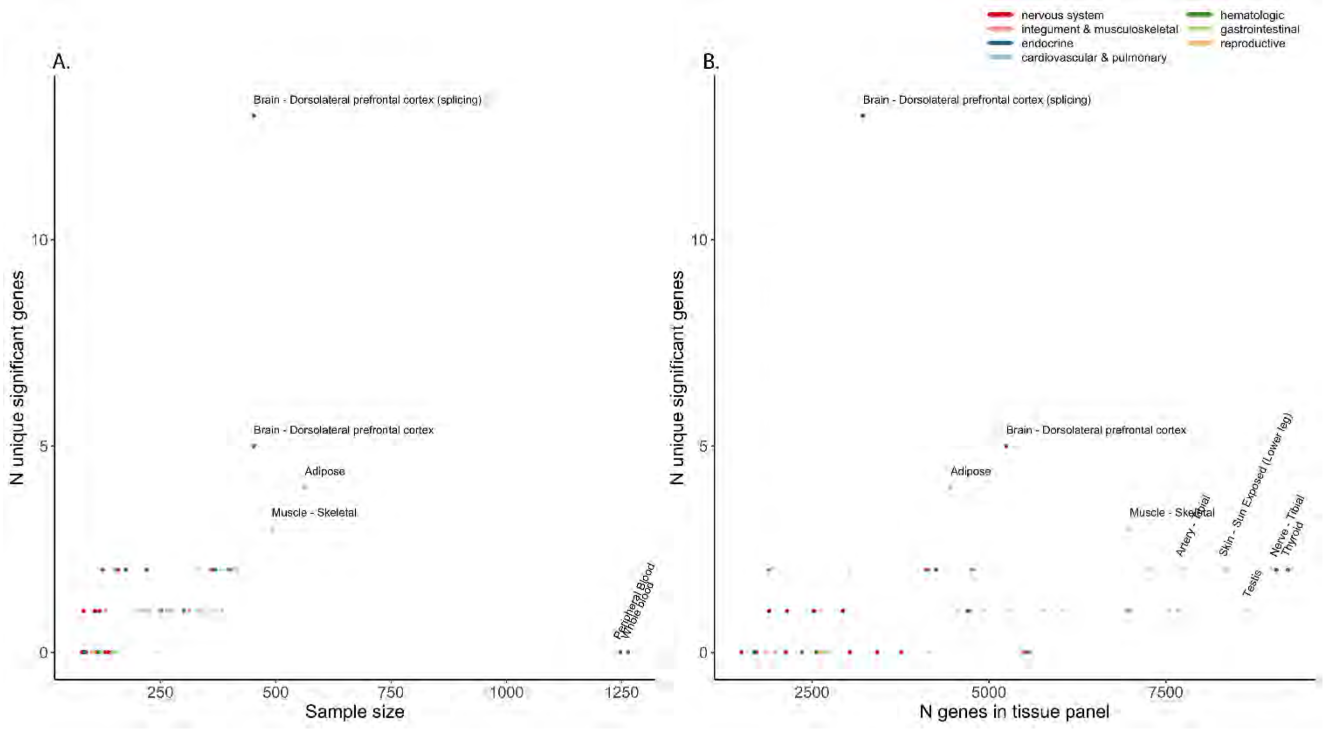


**Figure S3. Relationship between number of unique transcriptome-wide significant genes and sample size (A) and number of genes measured in the eQTL reference panel (B).**
eQTL; expression quantitative trait loci.

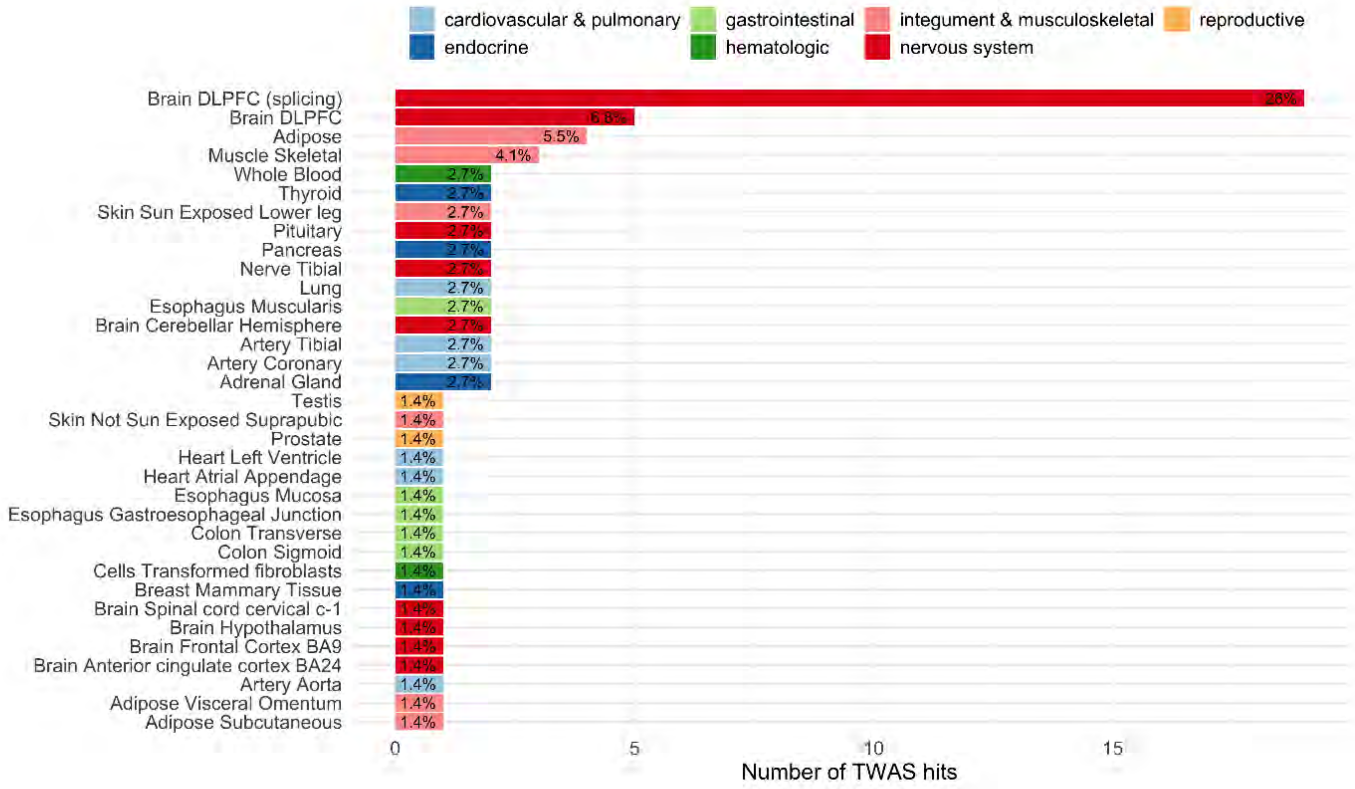


**Figure S4. Number of unique transcriptome-wide significant genes in FTD TWAS.**

FTD; frontotemporal dementia, TWAS; transcriptome-wide association study.

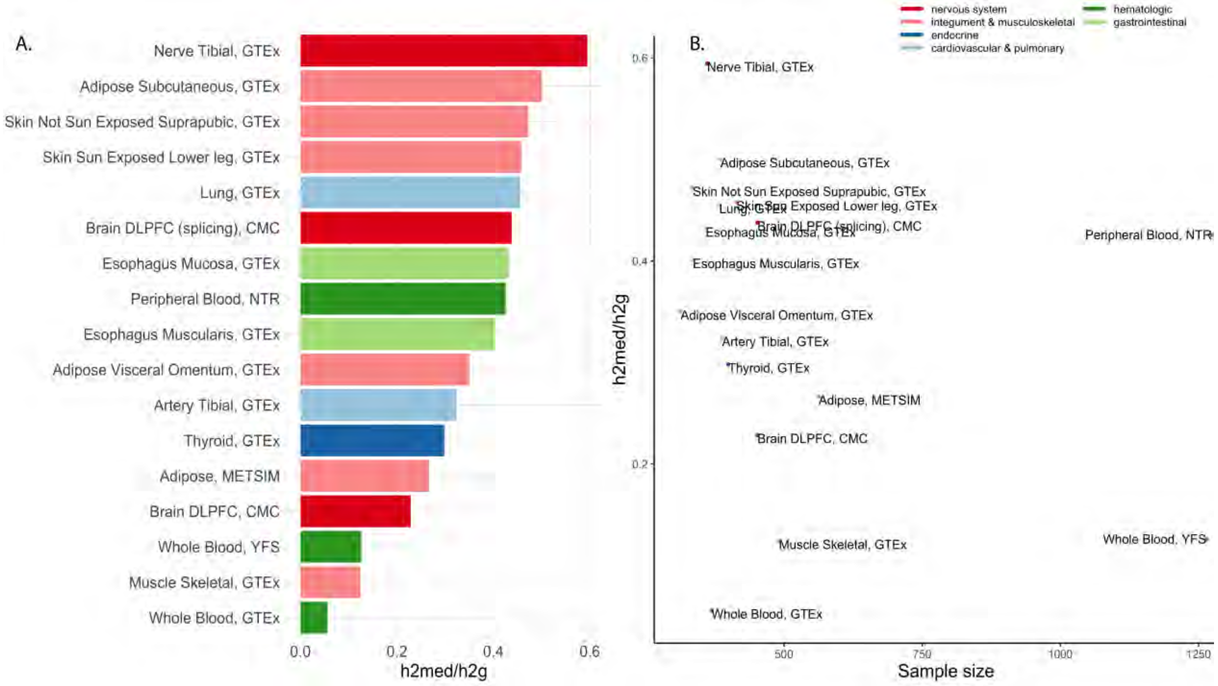


**Figure S5. Results MESC analysis on FTD for 17 eQTL reference panels with sample size n>300.** (A) Estimated proportion of heritability mediated by the cis-genetic component of gene expression (h2med/h2) per tissue type. (B) Relationship between heritability mediated by the cis-genetic component of gene expression (h2med/h2) and sample size. FTD; frontotemporal dementia, MESC; mediated expression score regression, eQTL; expression quantitative trait loci.

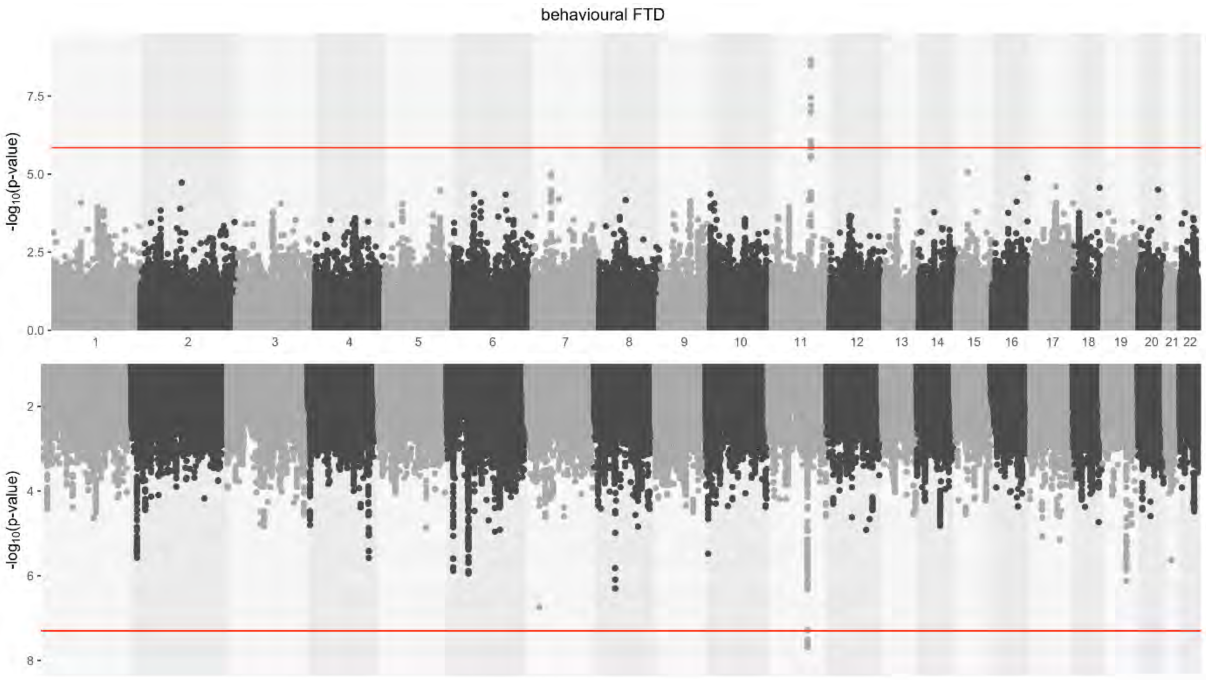


**Figure S6. Manhattan plot on bvFTD TWAS (top) and GWAS (bottom).**

bvFTD; behavioral frontotemporal dementia, TWAS; transcriptome-wide association study, GWAS: genome-wide association study.

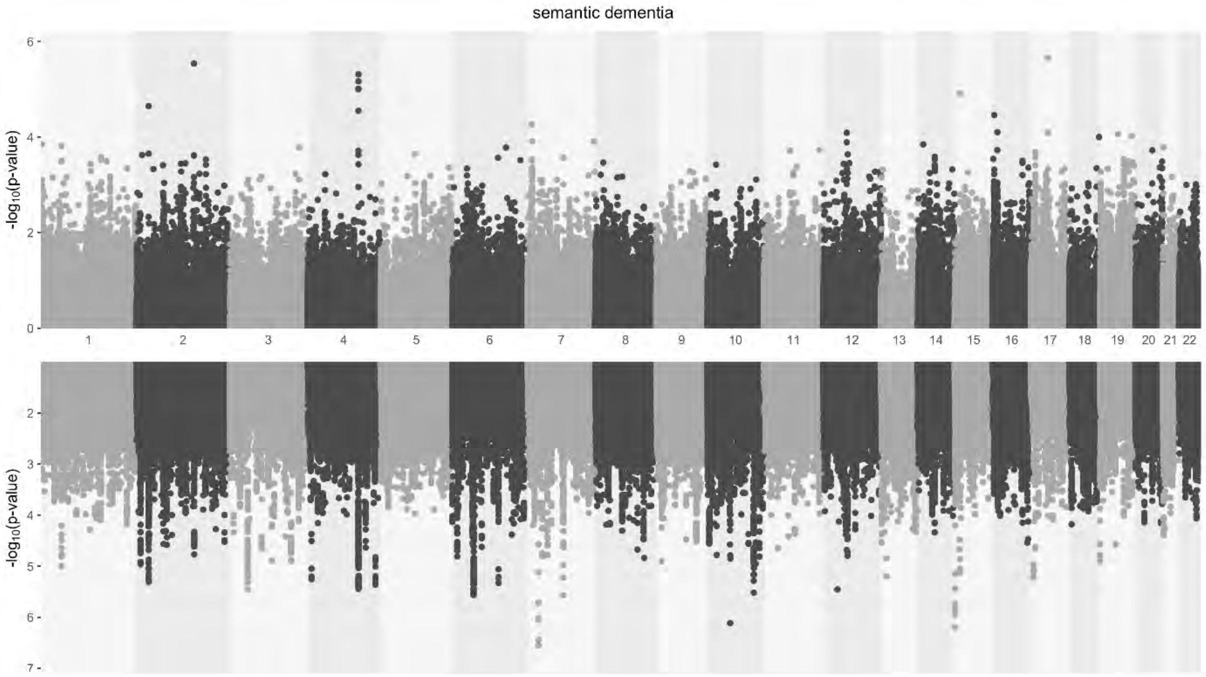


**Figure S7. Manhattan plot on SD TWAS (top) and GWAS (bottom).**

SD; semantic dementia, TWAS; transcriptome-wide association study, GWAS: genome-wide association study.

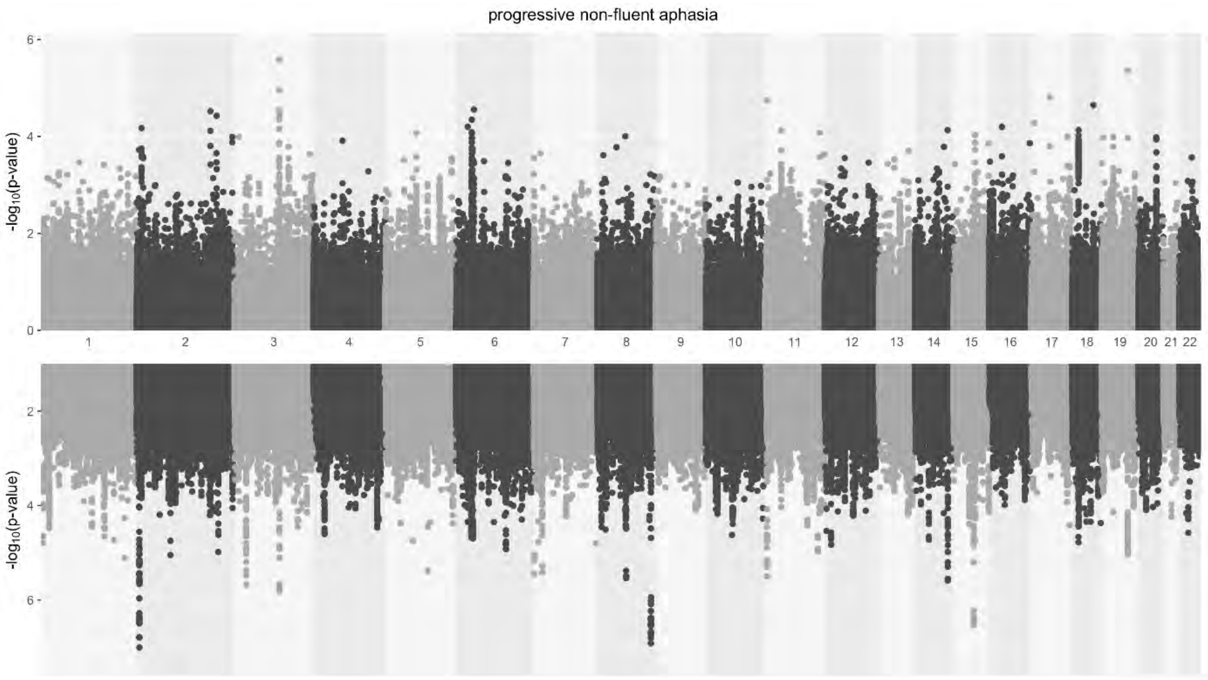


**Figure S8. Manhattan plot on PNFA TWAS (top) and GWAS (bottom).**

PNFA; progressive non-fluent dementia, TWAS; transcriptome-wide association study, GWAS: genome-wide association study.

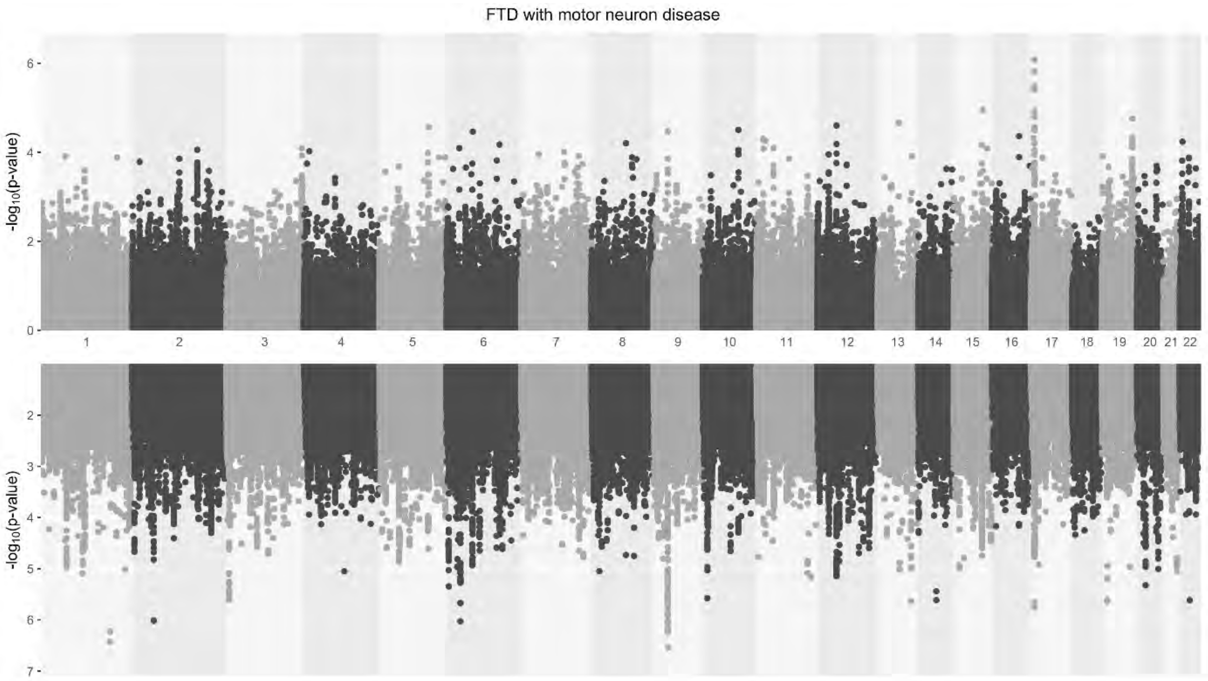


**Figure S9. Manhattan plot on FTD-MND TWAS (top) and GWAS (bottom).**

FTD-MND; frontotemporal dementia with motor neuron disease, TWAS; transcriptome-wide association study, GWAS: genome-wide association study.

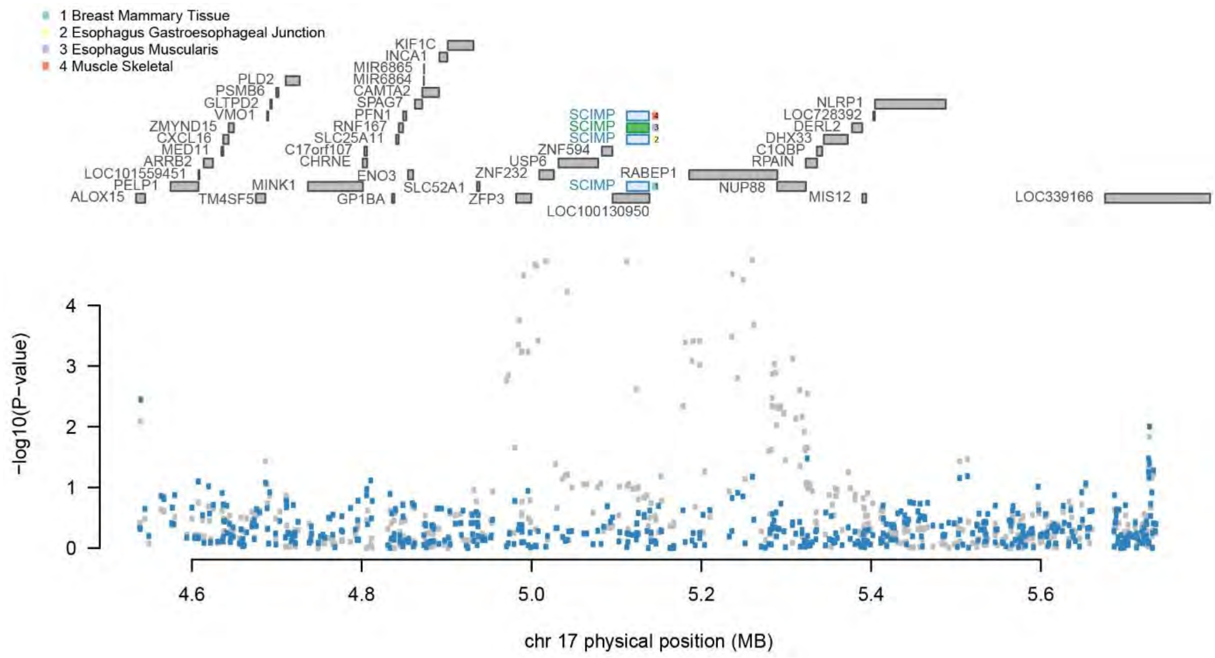


**Figure S10. Regional association plot of SCIMP for FTD with motor neuron disease.** The top panel shows all of the genes in the locus. The marginally TWAS associated genes are highlighted in blue, and those that are jointly significant (i.e., SCIMP in Esophagus Mascularis) highlighted in green. The bottom panel shows a Manhattan plot of the GWAS data before (gray) and after (blue) conditioning on the green genes. Most SNPs in this locus go from being genome-wide significant to non-significant after conditioning on the predicted expression of SCIMP.

FTD-MND; frontotemporal dementia with motor neuron disease, TWAS; transcriptome-wide association study, chr; chromosome, MB; megabase.

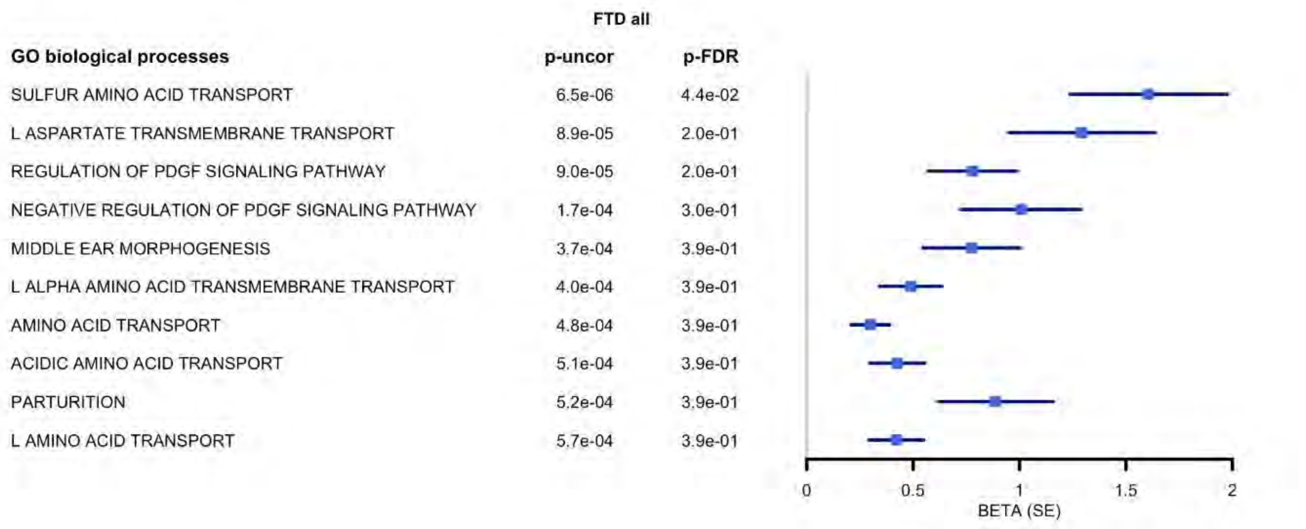


**Figure S11. Top 10 significant biological processes associated with TWAS FTD (all subtypes) results, including the MHC region.**

FTD; frontotemporal dementia, TWAS; transcriptome-wide association study, MHC; major histocompatibility complex, GO; gene ontology, uncor; uncorrected, FDR: false-discovery rate, SE; standard error.

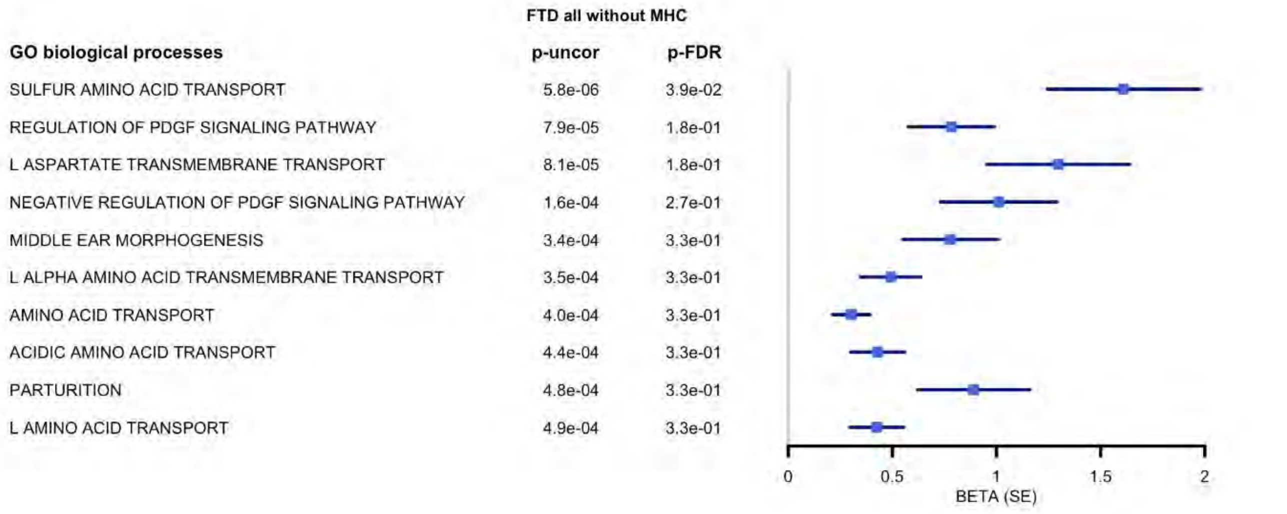


**Figure S12. Top 10 significant biological processes associated with TWAS FTD (all subtypes) results, excluding the MHC region**.

FTD; frontotemporal dementia, TWAS; transcriptome-wide association study, MHC; major histocompatibility complex, GO; gene ontology, uncor; uncorrected, FDR: false-discovery rate, SE; standard error.

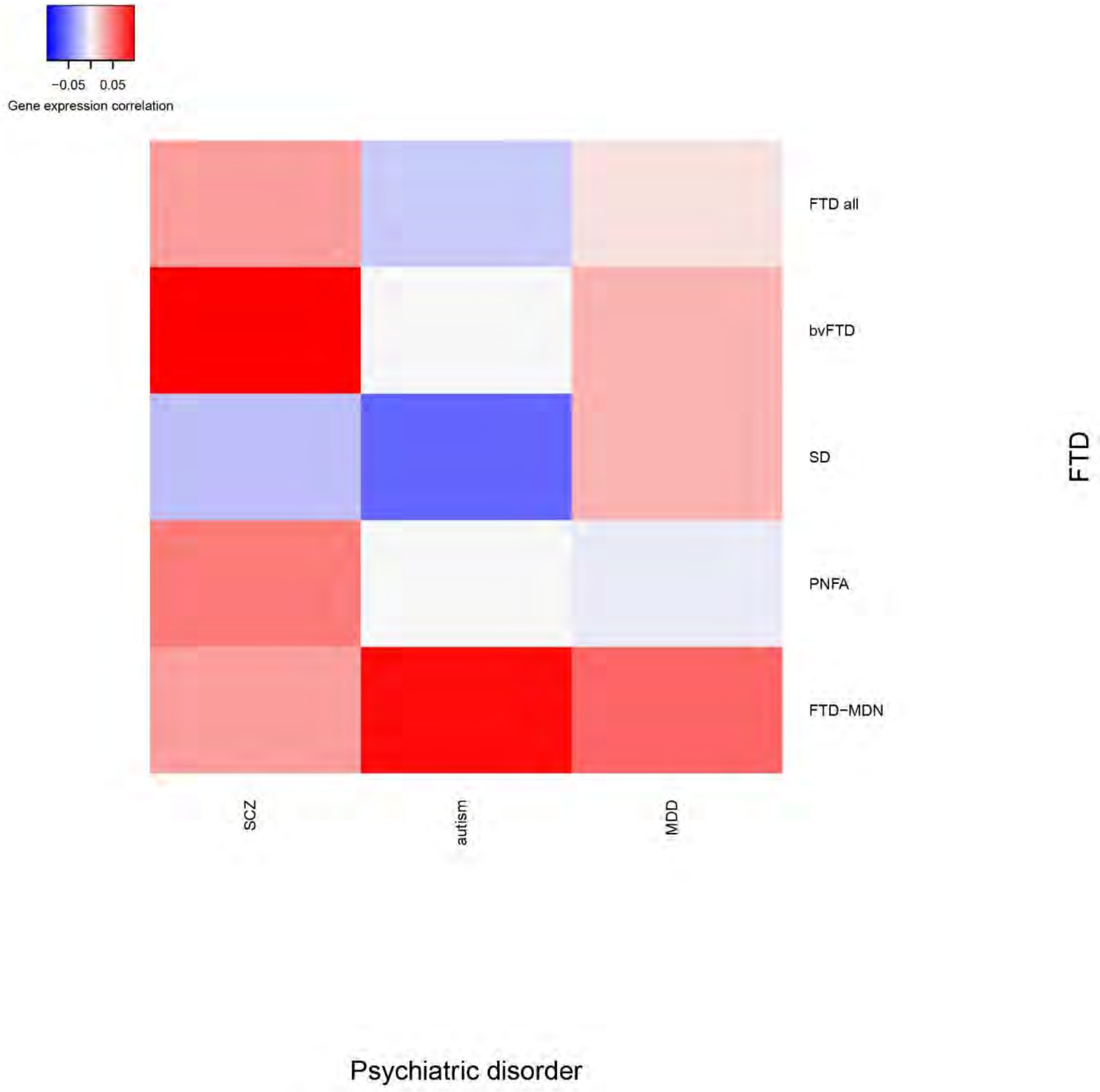


**Figure S13. Genetic correlation between gene expression of FTD and psychiatric disorders.**

FTD; frontotemporal dementia, bvFTD; behavioral variant FTD, SD; semantic dementia, PNFA; progressive non-fluent aphasia, FTD-MND; FTD with motor neuron disease, SCZ; schizophrenia, MDD; major depressive disorder.

#### Supplemental methods

##### Expression quantitative trait loci (eQTL) reference panels

The 53 eQTL reference panels included data on cardiovascular and pulmonary (n=6), nervous system (n=17), endocrine (n=5), reproductive (n=5), gastrointestinal (n=9), hematologic (n=5) and integument and musculoskeletal (n=6) tissues. The CMC includes RNA seq and RNA seq splicing data on the dorsolateral prefrontal cortex (DLPFC) from n=452 participants (n=254 schizophrenia, 52 bipolar, 146 controls, age of death 20-100y, 39.7% female). NTR includes RNA array data on peripheral blood from n=1,247 individuals (age 28–39y, 65.8% female). YFS contains whole blood gene expression data on n=1,264 individuals (age 34-49y, 55.9% female). METSIM contains adipose tissue gene expression data on n=562 males (age 45-74y). GTEx v7 includes gene expression data (RNA seq) from $n_{total}$=752 individuals (age of death 20-79y, 34.3% female) with different causes of death. GTEx contains gene expression on 48 tissue types ($n_{tissue}$=80-491), including cardiovascular & pulmonary (n=6), nervous system (n=15), endocrine (n=5), reproductive (n=5), gastrointestinal (n=9), hematologic (n=3) and integument & musculoskeletal (n=5) tissues (Table S25). All study subjects provided written informed consent.

##### TWAS colocalization analysis

COLOC estimates the posterior probability (PP) for: 0. No association with either FTD or gene expression (PP0), 1. Association with FTD only (PP1), 2. Association with gene expression only (PP2), 3. Association with FTD and gene expression from distinct causal variants (PP3) and 3. Shared causal variant (PP4). We defined colocalization evidence as supportive for a shared causal variant between the functional feature and FTD, when PP4>0.17 and PP4>PP3 as discussed in Gusev 2016 (Gusev *et al.*, 2016).

##### RHOGE analysis

Given the symptom overlap between FTD and major psychiatric disorders, we aimed to examine the genetic correlation between the predicted gene expression of FTD (and its clinical subtypes) and major psychiatric disorders, including schizophrenia, autism and major depressive disorder, using RHOGE software (https://github.com/bogdanlab/RHOGE)(Mancuso *et al.*, 2017). Given the complex LD structure of the MHC region, analyses were repeated with and without the MHC region. Results on fifteen phenotype pairs were corrected for multiple comparisons using a 5% FDR significance threshold.
